# sncRNAP: Prediction and profiling of full sncRNA repertoires from sRNAseq data

**DOI:** 10.1101/2023.05.23.541863

**Authors:** Hesham A. Y. Gibriel, Sharada Baindoor, Ruth S. Slack, Jochen H. M. Prehn

## Abstract

**Motivation:** Non-coding RNAs (ncRNAs), which include long non-coding RNAs (lncRNAs) and small non-coding RNAs (sncRNAs), have been shown to play essential roles in various biological processes. Over the past few years, a group of sncRNA identification tools have been developed but none has shown the capacity to fully profile and accurately identify those that are differentially expressed in control vs treated samples. Therefore, a tool that fully profiles and identifies differentially expressed sncRNAs in group comparisons is required.

**Results:** We developed sncRNAP, a Nextflow pipeline for the profiling and identification of differentially abundant sncRNAs from sRNAseq datasets. sncRNAP primary use case is the comparison of multiple small RNA-seq datasets belonging to two conditions such as the comparison of treatment (T) and control (C) cohorts. sncRNAP can be used to analyze human, mouse, and rat datasets. The pipeline carries out all the steps required to assess raw sequencing data, performs differential gene expression (DE) analysis, profiles sncRNAs in each sample, and outputs TXT, PDF, CSV, and interactive HTML files for the quality score and the top identified sncRNA candidates. We verified sncRNAP on publicly available sRNAseq datasets in chronic hepatitis-infected liver tissue and pancreatic ductal adenocarcinoma (PDAC) datasets. Our results support the identification of Val[C/A]AC in hepatitis patients and miR135b in PDAC as potential disease biomarkers. Furthermore, we applied sncRNAP on mouse samples from control and Opa1 mouse mutants and identified AspGTC, ValAAC, SerTGA, and AspGTC as the top DE tsRNAs. In addition, sncRNAP identified mmu-miR-136-5p, mmu-miR-10b-5p, mmu-miR-351-5p, and mmu-miR-6390 as the top DE miRNA candidates.

## Introduction

### Why small-non-coding RNAs (sncRNAs) are important?

Non-coding RNAs (ncRNAs), which include long non-coding RNAs (lncRNAs) and small non-coding RNAs (sncRNAs), have been previously viewed as molecules that are merely “junk”. However, recently, researchers have elucidated roles for ncRNAs in various biological processes. Such roles involve complex mechanisms that play crucial roles for example in development and carcinogenesis (Iyer *et al*., 2015; Anastasiadou *et al*., 2018; Yang *et al*., 2020; Wolin and Maquat, 2019). A major class of sncRNA are tRNAs that are known for their cloverleaf secondary structure and for delivering amino acids to the ribosome for protein synthesis (Krahn *et al*., 2020). During stress responses, tRNAs can be cleaved by ribonucleases including Dicer and angiogenin (ANG), yielding tRNA-derived small RNAs (tsRNAs) with new roles in gene expression, protein translation and cell signaling (Li *et al*., 2018). Such tsRNAs are classified into either tiRNAs, produced when tRNAs are cleaved in the anticodon loop by ANG, or tRNA-derived fragments (tRFs), which encompass the remainder of non-tiRNA tsRNAs (Li *et al*., 2018). In addition to tsRNAs, microRNAs (miRNAs) are a further, essential class of sncRNAs that are involved in post-transcriptional regulation of gene expression. There are also other classes of sncRNAs such as small nucleolar RNAs (snoRNAs), Y RNAs, Vault RNAs, and rRNAs that have been previously shown to play roles in different biological processes (Falaleeva and Stamm, 2013; Jackowiak *et al*., 2011; Chen and Heard, 2013). For example, snoRNA-93 have been shown to be involved in cell invasion in breast cancer (Patterson *et al*., 2017). All these previous examples elucidate the essential roles of sncRNAs in disease progression and highlight the need for further investigations to reveal additional sncRNA functions.

### Current sncRNA prediction tools are not comprehensive

As miRNAs are one of the major sncRNA classes, numerous tools have been previously developed to predict their abundance in sRNAseq datasets. One of these tools is miRDB, an online resource that utilizes MirTarget for miRNA target prediction in five species: human, mouse, rat, dog and chicken (Wong and Wang, 2015). Another tool is TargetScan, which identifies miRNA targets by identifying mRNAs with conserved complementarity to the seed (nucleotides 2-7) of the miRNA (Agarwal *et al*., 2015). Other tools such as miRgo combine multiple tools into integrated systems to develop a novel prediction system (Chu *et al*., 2020). Recently, smrnaseq, a nextflow pipeline for the identification of differentially expressed miRNAs from sRNAseq datasets has been developed (Pantano *et al*., 2022). The pipeline reports novel miRNAs in sRNAseq samples and produces reports for reads, alignments, and expression results (Pantano *et al*., 2022). In addition to miRNA identification, tools have been developed previously to facilitate the identification of tsRNAs from sRNAseq datasets. MINTmap is one such tool that can identify abundant tRFs in RNAseq datasets (Loher *et al*., 2017). The tool generates detailed tRF expression profile through an exhaustive genome-wide search (Loher *et al*., 2017). SPORTS1.0 is a pipeline for identifying short RNAs from 68 species (Shi *et al*., 2018). tDRmapper is a standalone command line interface (CLI) pipeline that reports reads from human and mouse samples between 14 and 40 nucleotides in length (Selitsky and Sethupathy, 2015). Although the previous miRNA and tsRNA tools can facilitate the identification of miRNAs and tRNAs in sRNAseq samples, they either lack a downloadable version, limited to a single sRNAseq datasets, or lack the comparative analysis of multiple replicates between different conditions.

To avoid the limitations in the previous miRNA and tsRNA tools and to build a tool that identifies miRNAs as well as other ncRNAs, tsRNAsearch was recently introduced (Donovan *et al*., 2021). tsRNAsearch is a Nextflow pipeline for the identification of differentially expressed tRNA fragments and other non-coding RNAs between two groups, such as control and treatment groups, from sRNAseq datasets (Donovan *et al*., 2021). tsRNAsearch reports tRNAs and ncRNAs based on a combined score that is made up of four novel methods (Donovan *et al*., 2021). However, tsRNAsearch uses miRNA hairpin annotations, which is insufficient for profiling the full length miRNA, and the combined score method adds a layer of ambiguity to the pipeline, as often the top identified candidates based on the combined score show low expression pattern and *vice versa*.

### Difference between sncRNAP and previous pipelines

All of the above pipelines for sncRNA predictions have limitations including: the previous pipelines, except tsRNAsearch, cannot be run for group and paired analysis, 2) the previous pipelines, except tsRNAsearch, cannot be run to profile more than one class of sncRNAs, 3) lack of paired analysis, 4) high computational power and low speed performance, and 5) improper adapter sequence detection which yields erroneous sncRNA identification (Fig. 1). As a result, sncRNA prediction will be compromised unless researchers run all these tools together which is time consuming and may lead to conflicts in the outputs.

**Fig. 1.**
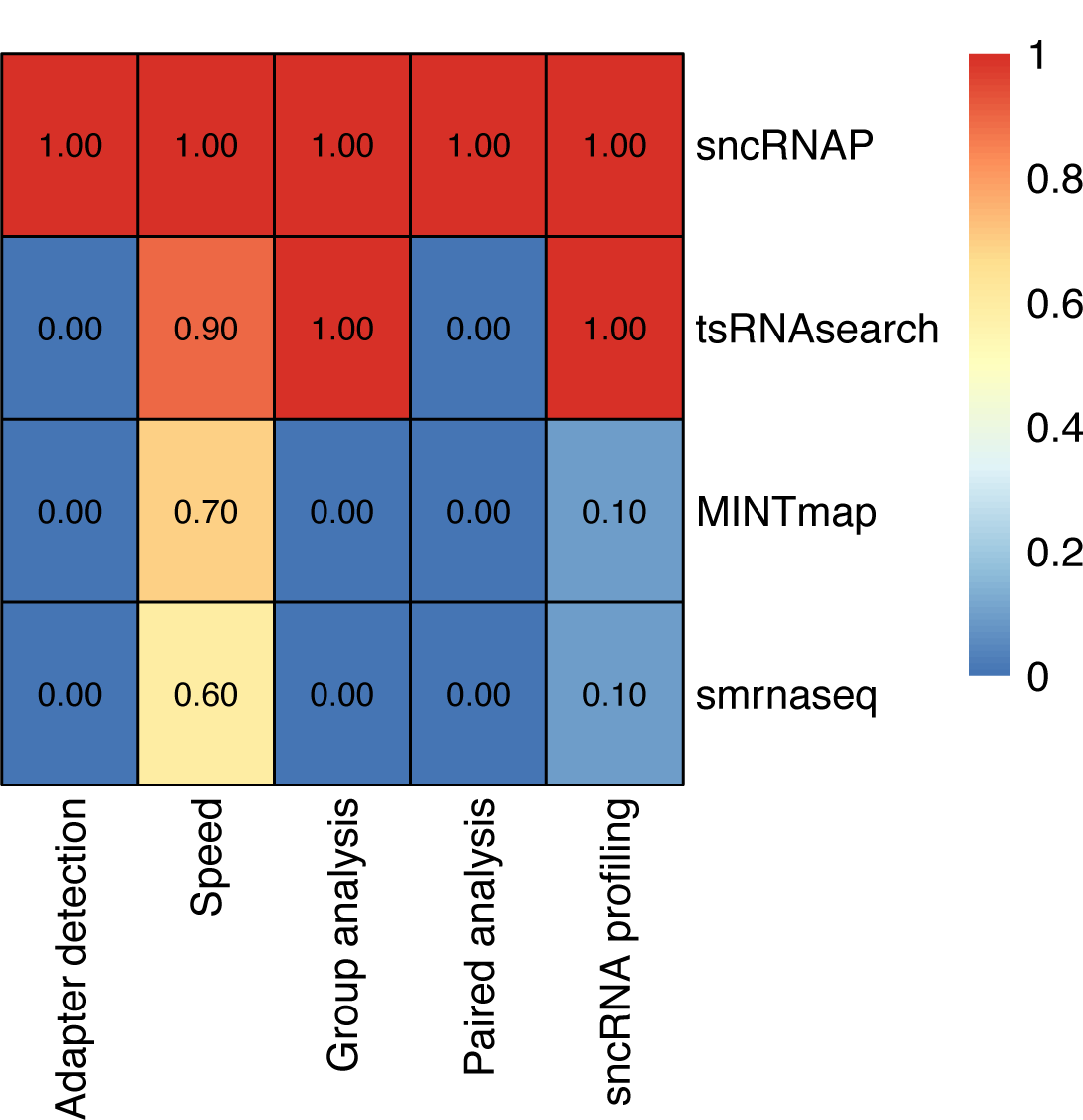
Comparison between sncRNAP and previously known pipelines for sncRNA identification. sncRNAP is advantageous over all these pipelines as it can detect adapter sequence from sRNAseq datasets as well as having the capacity to perform faster, fully profile sncRNAs, and apply group and paired analysis.

Here, we developed sncRNAP, a Nextflow pipeline for the profiling and identification of differentially abundant sncRNAs from sRNAseq datasets. sncRNAP primary use case is the comparison of multiple small RNA-seq datasets belonging to two conditions such as the comparison of treatment (T) and control (C) cohorts. sncRNAP can be used to analyze human, mouse, and rat datasets. By default, sncRNAP carries out all the steps required to assess raw sequencing data and output results such as quality control, pre-processing, alignment, and post-processing (Fig. 2). Additionally, sncRNAP profiles sncRNAs in each sample. In this manner, sncRNAP shows the length distribution, sncRNA counts per sample, overlapping sncRNA in each sample, sncRNA expression per sample and per sncRNA class, volcano plots for each sncRNA class and heatmap for the top differentially expressed sncRNAs. Lastly, sncRNAP reports quality scores as well as the fasta sequences for the top identified candidates in the experiment.

**Fig. 2.**
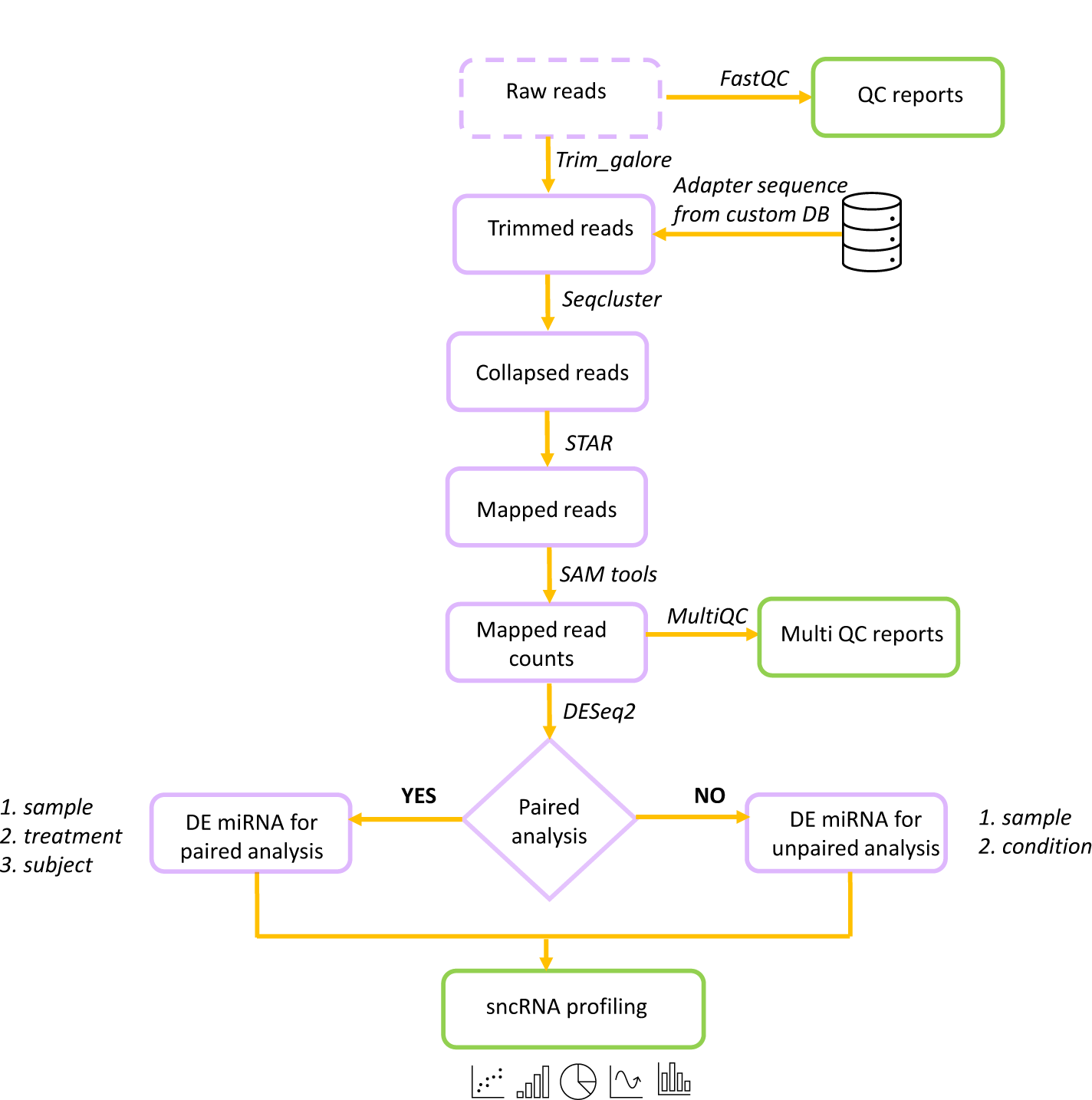
sncRNAP pipeline workflow. sncRNAP is a nextflow pipeline that identifies differentially expressed sncRNAs from group analysis such as C vs T in sRNAseq datasets. The main steps include pre-processing and it ends with identification of differentially expressed sncRNAs from different groups whether these conditions are considered as paired or not.

## Methods

### 1.1 Pipeline overview

sncRNAP is a Nextflow pipeline (di Tommaso *et al*., 2017) with package and environment control taken care of by conda. This allows for increased potential across a variety of different Unix systems. The pipeline can be run using the following command:

‘nextflow run sncRNAP –input_dir data/--output_dir./Results--genome human--layout layout.csv-–paired_samples FALSE’ A description of the steps in the pipeline are as follows (Fig. 2):

1. Quality check reads (FastQC) (Andrews, 2010)
2. index db for sncRNA database (STAR) (Dobin *et al*., 2013)
3. Adapter sequence identification (get_adapter.py)
4. Adapter trimming (trim_galore) (Andrews, 2015)
5. Read collapse (seqcluster) (Pantano *et al*.)
6. Alignment against sncRNA database (STAR (Dobin *et al*., 2013)
7. Differential expression analysis (DESeq2) (Love *et al*., 2014)
8. sncRNA profiling (sncRNAP_profyling.py)
9. sncRNA quality control (mirtrace) (Kang *et al*., 2018) and QC for raw read, alignment, and expression reports (MultiQC) (Ewels *et al*., 2016)

### 1.2 sncRNAP output

sncRNAP outputs TXT, PDF, CSV, and interactive HTML files on the quality score and the top identified sncRNA candidates. The PDF reports the length distribution, sncRNA counts per sample, overlapping sncRNA in each sample, sncRNA expression per sample and per sncRNA class, volcano plots for each sncRNA class, sncRNA abundance, and heatmap for the top differentially expressed sncRNAs. The HTML files provide general stats on percentage of mapped reads, featurecounts, number of reads, number of reads removed by Cutadapt, fastqc reports, read length distribution, percentage of sncRNA reads, and RNA type. The TXT file reports the top identified sncRNA fasta sequences and the CSV file reports the log2fc, pvlaue, padj, and RPKM values for the top sncRNA candidates for each sample. Finally, the pipeline run information is generated by Nextflow for each step of sncRNAP.

### 1.3 Pipeline default parameters

The minimum alignment length is 16 nucleotides, which has been chosen to avoid false mapping to genomic sequences identical to sncRNA fragments under 15 nucleotides in length. By default half of the available free CPU resources are set to be used for the pipeline execution and this can be overridden when executing the pipeline to use less or more resources. Differential expression threshold is ((log2FoldChange > 1 or log2FoldChange <-1) & padj < 0.005). If the option for paired group analysis is not specified the pipeline conducts unpaired group analysis by default. A detailed description of sncRNAP and the available parameters are available at https://github.com/hesham123457887/sncRNAP. Packages used include: Nextflow (di Tommaso *et al*., 2017), ggplot2 (Villanueva and Chen, 2019), gplots (Warnes et al., 2013), ggrepel (Slowikowski, 2019), DESeq2 (Love *et al*., 2014), Python (Rossum, 1993), R (Team, 2017), reshape2(Wickham, 2007) and dplyr(Wickham et. al., 2022).

### 1.4 sncRNAP databases

sncRNAP currently includes three datatabses from human, mouse, and rat. These databases were retrieved from tsRNAsearch (Donovan *et al*., 2021) with the modification, addition, and deletion of previously existing sncRNA ids. In this manner, all miRNA sequences retrieved from tsRNAsearch databases were removed, as those are only hairpin miRNAs sequences, and replaced with mature miRNA sequences from miRbase (Kozomara *et al*., 2018). Additionally, to distinguish sncRNAs classes, ids have been modified to include gene ids from Ensembl followed by the ncRNA class.

### 1.5 sncRNAP on simulated data

We generated a simulated dataset to detect the sensitivity, specificity and precision of the sncRNAP pipeline (Supplementary figure 1). We combined the sncRNAP database sequences with sequences from exomic coding sequence (cds) from Ensembl that are considered as negative controls. STAR was used to align the simulated reads to the sncRNAP STAR database and the whole processes were repeated 10 times for each species. Subsequently, true positives (TP), true negatives (TN), false positives (FP), and false negatives (FN) were scored and the average sensitivity, specificity and precision were calculated (Table 1).

**Table 1.**
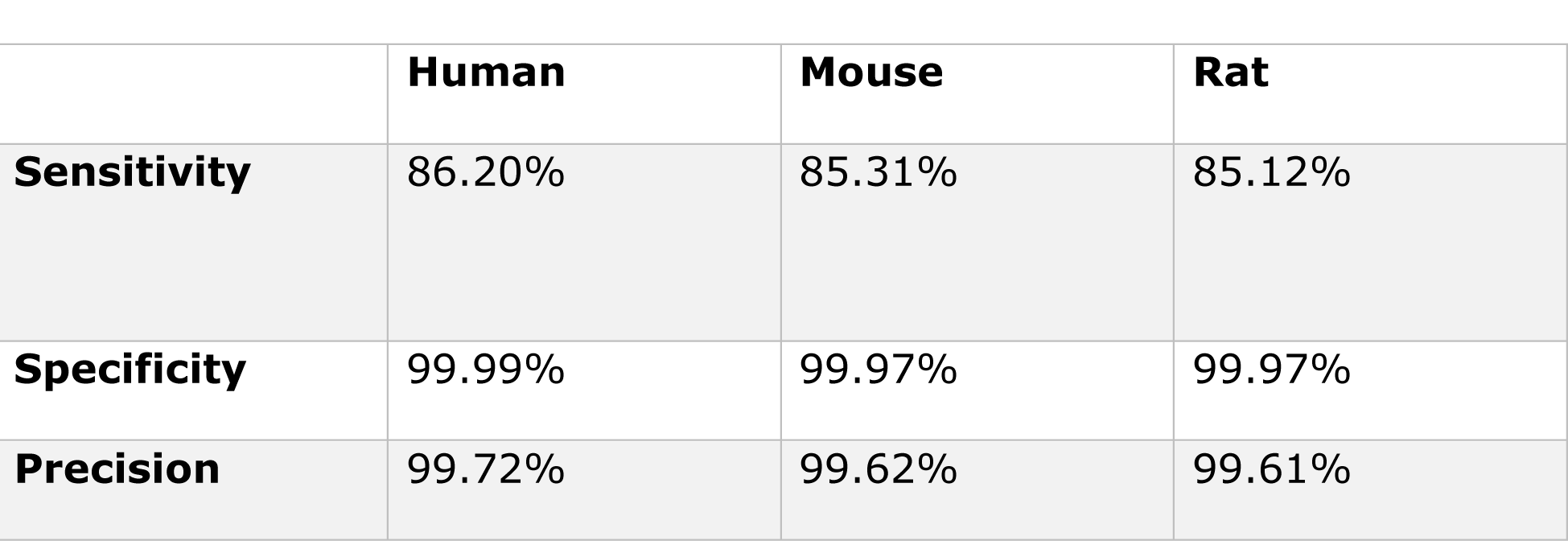
sncRNAP sensitivity, specificity and precision on simulated datasets.

|  | Human | Mouse | Rat |
| --- | --- | --- | --- |
| **Sensitivity** | 86.20% | 85.31% | 85.12% |
| **Specificity** | 99.99% | 99.97% | 99.97% |
| **Precision** | 99.72% | 99.62% | 99.61% |

### 1.6 Opa1 dataset

sncRNAP was applied to sRNAseq dataset generated from six samples from control and Opa1 mouse mutants collected at day 3 with NEBNext small RNA library prep. The dataset contained two groups (day 3 and day 5) six samples in each group. These groups included wild-type vs Opa1 knock-down with no AlkB pre-treatment.

### 2.7. Public datasets

The ncRNA-seq data (GEO: GSE57381) analyzed in the study contained five groups, with four samples in each group (Selitsky *et al*., 2015). These groups included control liver tissue, liver tissue from patients with hepatitis B and associated cancer, hepatocellular carcinoma (HCC) tissue from patients with hepatitis B, liver tissue from patients with hepatitis C and associated cancer, and HCC tissue from patients with hepatitis C (Selitsky *et al*., 2015). The authors created these data to identify differentially expressed tsRNAs, and therefore we ran sncRNAP on this dataset.

In addition to the previous dataset, we ran sncRNAP to a paired dataset containing seven pancreatic ductal adenocarcinoma (PDAC) samples and seven healthy control samples (GSE125538) (Yang *et al*., 2019). The data was originally generated to characterize miRNAs in PDAC tumors, and therefore considered as a proper validation of miRNAs (Yang *et al*., 2019).

### 2.8 Comparison of sncRNA identification tools

We compared sncRNAP to other tools to determine the pipeline’s ability to detect sncRNAs. A comparison was conducted between sncRNAP, smRNAseq (Pantano *et al*., 2022), and tsRNAsearch (Donovan *et al*., 2021) on FASTQ files retrieved from the GSE125538 database (Yang *et al*., 2019). Reads aligned to sncRNAs from each tool were pooled and the Pearson correlation coefficients between the sncRNA read count for each tool was calculated to determine how sncRNA tools corelate with each other.

## 3 Results

### 3.1 High sensitivity, specificity, and precision scores on simulated datasets

We used sncRNAP to measure the sensitivity, specificity and precision on human, mouse, and rate simulated datasets. For all datasets, sncRNAP showed high sensitivity (∼85%), high specificity (∼99%), and high precision (∼99%) (Table 1). Specifically, the human datasets scored the highest of these tests (Table 1).

### 3.2 sncRNAP is comparable with previous tools and with more advantages

sncRNAP, smRNAseq (Pantano *et al*., 2022), and tsRNAsearch (Donovan *et al*., 2021) were applied on FASTQ files retrieved from the GSE125538 PDAC database (Yang *et al*., 2019). We observed that sncRNAP and smRNAseq were highly correlated, with an average Pearson’s *r* = 0.85, while the comparison between sncRNAP and tsRNAsearch scored an average Pearson’s *r* = 0.61.

In addition to sncRNAP ability to accurately map reads to sncRNA databases, sncRNAP has more advantages that are not present in the previously compared tools. In this manner, sncRNAP can precisely identify adapter sequence in fastq files (Fig. 1), while the previous tools rely on automatic adapter sequence identification without searching if the adapter sequence is present in the fastq read. This is a crucial step as if the correct adapter sequence is not identified and removed from a fastq file, it can lead to incorrect sequence alignment in the downstream analysis. Moreover, sncRNAP running time is less than the other tools (Fig. 1, Supplementary figure 2), and is the only tool that performs paired analysis (Fig. 1). Lastly, similar to tsRNAsearch, sncRNAP profiles the full sncRNA repertoire (Fig. 1), with the advantage of accurately profiling miRNAs as it uses the mature rather than the hairpin miRNA sequences.

### 3.3 sncRNAP confirmed the identification of Val[C/A]AC and identified other sncRNAs candidates in hepatitis patients

sncRNAP was applied to a dataset containing samples from control patients, patients with hepatitis B and an associated cancer and hepatitis C with an associated cancer. The pipeline identified ∼18,461,476 reads per sample, of which ∼17,321,251 are QC-passed reads (Supplementary figure 7). The identified adapter sequence was ‘TGGAATTCTCGGGTGCCAAGG’ which has been identified in more than 88.6% of the sequenced reads (Supplementary figure 7). Read length distribution varied across samples, which ranged from 20-25 bp and 30-35 bp (Supplementary figure 3A).

The original authors found that tsRNAs are the most abundant small ncRNAs in hepatitis B and C infected liver tissue, and that tsRNA abundances are altered in HCC (Selitsky *et al*., 2015). We found that the majority of reads are originating from miRNAs followed by tRNAs, and rRNAs (Supplementary figure 3B). sncRNA counts were variable across samples, with misc_RNAs, miRNAs, snoRNAs, and tRNAs are the most present sncRNAs (Supplementary figure 3C).

sncRNAP subsequently profiled differentially expressed (DE) sncRNAs in all samples and identified 141 DE sncRNAs (Supplementary figure 3D). Of these, 94 (66.7%) tRNAs, 4 (2.8%) snoRNAs, 2 (1.4%) snRNAs, 30 (21.3%) misc_RNAs, and (11 7.8%) miRNAs were identified (Supplementary figure 3E). Similar to Selitsky and Sethupathy (Selitsky *et al*., 2015), we observed that 5-Gly[G/C]CC and 5-Val[C/A]AC are one of the most abundant tRNA fragments (Supplementary figure 3F). In addition to Val, we observed that Lys[C/T]TT, Gly[G/C]CC, CysGCA, Leu[C/A]AG, and Glu[C/T]TC are also one of the most abundant tsRNAs (Supplementary figure 3F).

DE analysis revealed ValCAC, ValAAC, LysCTT, CysGCA, GlyCCC, and GlyGCC as the top candidates for tsRNAs (Supplementary figure 4). Other identified DE sncRNA include ENSG00000206585 and ENSG00000206702 for snRNAs, ENSG00000240877 and ENSG00000244112 for misc_RNAs, ENSG00000202093, ENSG00000238649 for snoRNAs, and hsa-miR-224-5p, hsa-miR-182-5p, hsa-miR-217-5p, hsa-miR-216a-3p, and hsa-miR-4508 for miRNAs (Supplementary figure 4).

### 3.4 sncRNAP confirmed the identification of miR135b and identified other sncRNAs candidates in PDAC

sncRNAP was used to analyze 14 paired pancreatic samples (seven PDAC and seven adjacent normal tissue samples) (Yang *et al*., 2019). The pipeline identified ∼16,631,242 reads per sample, of which ∼15,228,319 are QC-passed reads (Supplementary figure 8). The identified adapter sequence was ‘TGGAATTCTCGGGTGCCAAGG’ which has been identified in more than 87.5% of the sequenced reads (Supplementary figure 8). Read length distribution varied across samples, which ranged from 20-25 bp and 30-35 bp (Fig. 5A), and the majority of reads are originating from miRNAs followed by tRNAs, and rRNAs (Supplementary figure 5B). sncRNA counts were variable across samples, with misc_RNAs, miRNAs, snoRNAs, and tRNAs are the most present sncRNAs (Supplementary figure 5C).

DE analysis revealed 19 DE sncRNAs (Supplementary figure 5D). Of these, one (5.3%) tRNAs, one (5.3%) misc_RNA, two (10.5%) rRNA, and 15 (78.9%) miRNAs were identified (Supplementary figure 5E). In the original study, the authors identified miR135b as significantly increased in PDAC samples compared to normal controls using a student’s t-test (P values < 0.05) on filtered normalized miRNA counts (Yang *et al*., 2019). Similarly, we observed that hsa-miR-135b-3p is one of the top DE miRNA candidates (Supplementary figure 6). In addition, sncRNAP identified hsa-miR-147b-3p, hsa-miR-210-3p, hsa-miR-552-3p, hsa-miR-31-5p, and hsa-miR-216b-3p as top DE candidates (Supplementary figure 6). Other identified DE sncRNA candidates include ArgTCT for tsRNAs, ENSG00000202111 for misc_RNAs, and ENSG00000276871 and ENSG00000283274 for rRNAs (Supplementary figure 6).

### 3.5 mmu-miR-136-5p and mmu-miR-10b-5p as well as other sncRNAs are differentially expressed in OPA1 mutants

For a further application of the pipeline, we isolated sncRNAs from six samples from control and Opa1 mouse mutants. sncRNAP was applied to this dataset and processed ∼13,231,112 reads per sample, of which ∼12,131,967 were QC-filtered (Supplementary figure 9). The identified adapter sequence was ‘AGATCGGAAGAGCACACGTCTGAAC’ which has been identified in more than 87.1% of the sequenced reads (Supplementary figure 9). Read length distribution varied across samples, the majority of which ranged from 20-25 bp and with less ranged from 30-35 bp (Fig. 3A). The majority of reads are originating from miRNAs followed by rRNA, and tRNAs (Fig. 3B), and the majority of sncRNA counts were miRNAs, snoRNAs, snRNAs, misc_RNAs, and tRNAs (Fig. 3C).

**Fig. 3.**
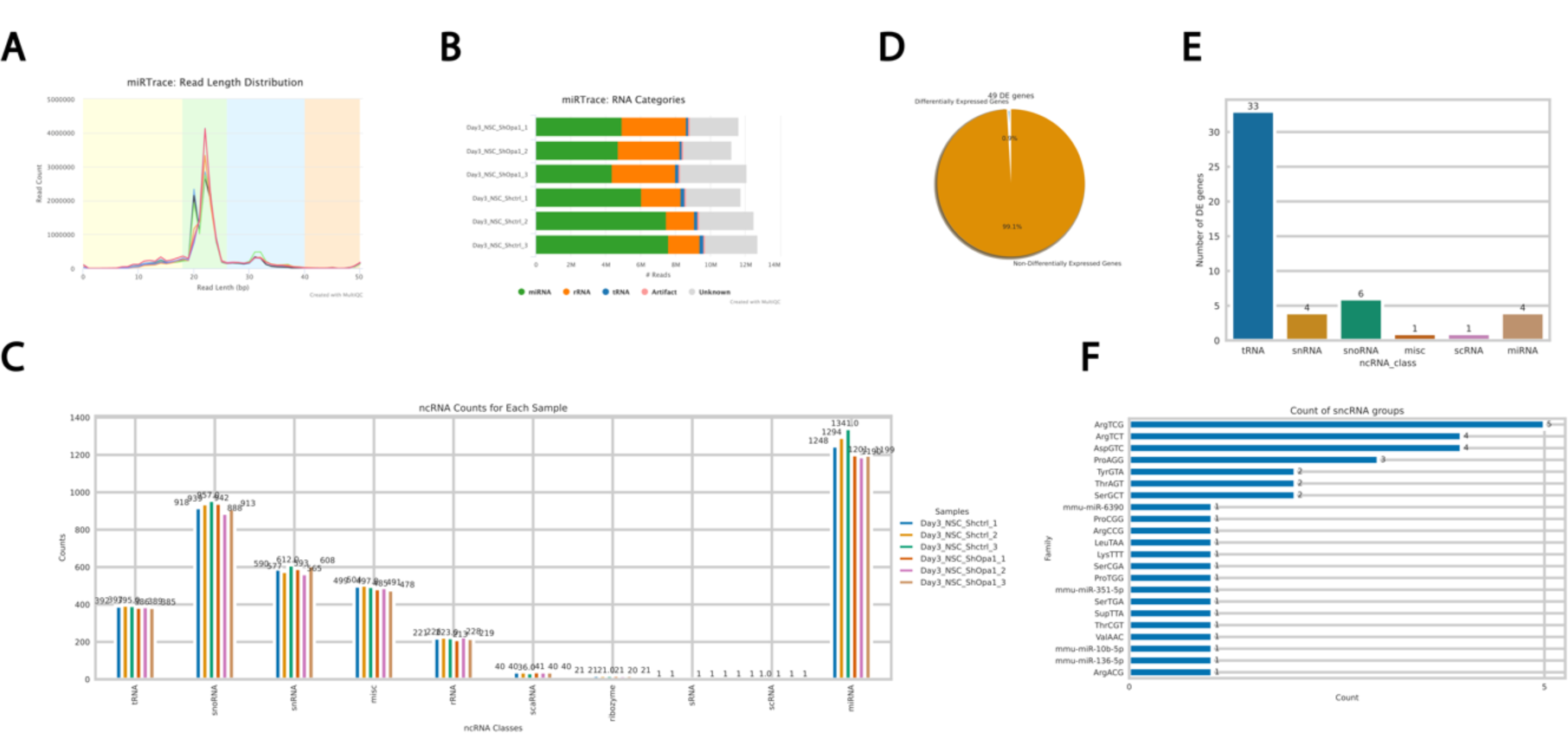
sncRNAP identified differentially expressed sncRNAs in OPA1 mutants. **A)** Read length distribution in bp of sncRNA reads. The majority of reads are between 20-25 and 30-35 bp. **B)** Percentages of RNA categories (miRNA, rRNA, and tRNA) for each sample. **C)** Counts of sncRNA classes for all samples. **D)** Pie chart showing the percentage for differentially and non-differentially expressed sncRNAs. **E)** Barplot showing the number of DE ((log2FoldChange > 1 or log2FoldChange < −1) & padj < 0.005) genes for each sncRNA class. **F)** Counts of the identified DE tRNA and miRNA families.

sncRNAP subsequently profiled DE sncRNAs in all samples and identified 49 DE sncRNAs (Fig. 3D). Of these, 33 (67.3%) tRNAs, four (8.2%) snRNAs, six (12.2%) snoRNAs, one (2%) misc_RNAs, one scRNA (2%), and four (8.2%) miRNAs (Fig. 3E). The most abundant tRNA fragments were ArgTC[G/T], AspGTC, ProAGG, TyrGTA, ThrAGT, and SerGCT (Fig. 3F), and The top DE candidates were AspGTC, ValAAC, SerTGA, and AspGTC (Fig. 4). In addition to tsRNAs, sncRNAP identified mmu-miR-136-5p, mmu-miR-10b-5p, mmu-miR-351-5p, and mmu-miR-6390 as the top DE miRNA candidates (Fig. 4). Other identified sncRNAs include: ENSMUSG00000077611 and ENSMUSG00000089414 for snoRNA, ENSMUSG00000065176 and ENSMUSG00000064851 for snRNA, and ENSMUSG00000098994 for misc_RNA, and ENSMUSG00000109440 for scRNA (Fig. 4).

**Fig. 4.**
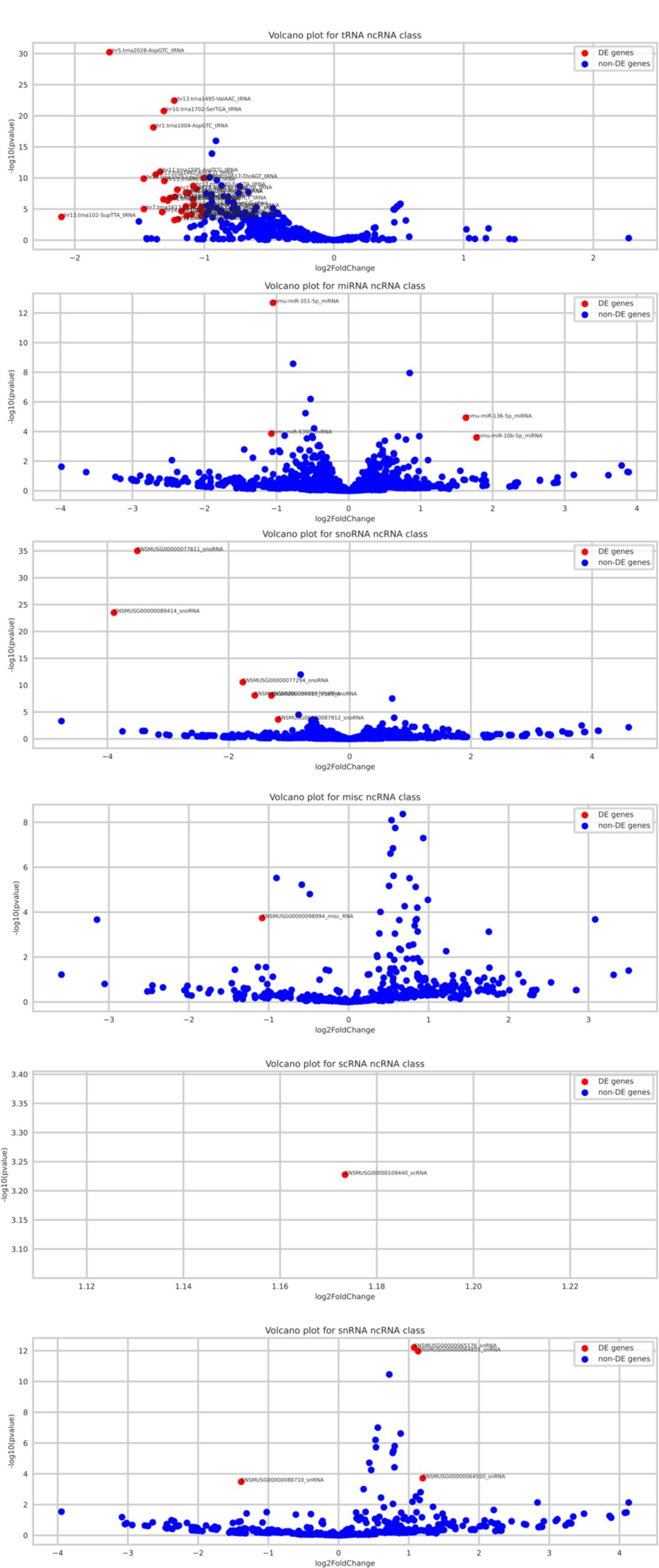
Volcano plots for the top identified differentially expressed sncRNAs in OPA1 mutants.

## Discussion

The availability of large-scale sequencing data from various sources has led to a plethora of sequencing data derived from different species, tissues, cell lines, and treatments. This is particularly relevant for sequencing data derived from sncRNAs, which yielded a significant amount of sncRNAs, such as miRNA and tRNAs, with the aim of identifying differentially expressed sncRNAs. While several tools have been developed for sncRNA identification, they have not been able to fully profile and accurately identify differentially expressed sncRNAs in group comparisons. Consequently, valuable sequencing information are often buried within the vast sequencing databases, making it challenging to associate sncRNAs to biological questions, for example which sncRNA can be used as a disease biomarker. Here, we developed sncRNAP, a Nextflow command line interface tool for the detection, quantification, and profiling of sncRNA fragments in sncRNAseq datasets. By utilizing sncRNAP, researchers can obtain useful information from their sncRNAseq datasets by accurately identifying differentially expressed sncRNAs in various species, tissues, cell lines, and treatments, providing a valuable resource for gaining insights into the roles of sncRNAs in diverse biological processes.

To verify sncRNAP results, we compared liver tissue from healthy patients with that of patients presenting with chronic hepatitis and an associated cancer (Selitsky *et al*., 2015). Similar to the study results, we found that tsRNAs are the most abundant small ncRNAs in hepatitis B and C infected liver tissue, especially those deriving from Valine and Lysine tRNAs, and also verified that 5-Gly[G/C]CC and 5-Val[C/A]AC are one of the most abundant tRNA fragments. In addition to Val, we observed that Lys[C/T]TT, Gly[G/C]CC, CysGCA, Leu[C/A]AG, and Glu[C/T]TC are also one of the most abundant tsRNAs. The pipeline also identified other sncRNAs that could be playing roles in disease progression such as hsa-miR-224-5p, hsa-miR-182-5p, hsa-miR-217-5p, hsa-miR-216a-3p, and hsa-miR-4508. We also verified sncRNAP on PDAC dataset and verified the finding that hsa-miR-135b-3p is one of the top DE miRNA candidates. The pipeline also identified hsa-miR-147b-3p, hsa-miR-210-3p, hsa-miR-552-3p, hsa-miR-31-5p, and hsa-miR-216b-3p as top DE miRNA candidates and ArgTCT as top tRNA candidates. Thus, sncRNAP successfully identified previously verified tRNA and miRNA biomarkers and identified additional sncRNAs that will be interesting to see if they have roles in disease progression.

Overall, sncRNAP is a novel Nextflow pipeline for the identification and profiling of sncRNAs in sncRNAseq datasets. The ideal use-case for this tool is to fully profile DE sncRNAs in the comparison between two cohorts of sncRNAseq datasets such as treatment and control groups from human, mouse and rat datasets.

## Acknowledgements

The authors acknowledge the grants supported to JHMP from Science Foundation Ireland (SFI) under the JPND program ‘RNA-NEURO’ [17/JPND/3455], the SFI Research Centre for Chronic and Rare Neurological Diseases ‘FutureNeuro’ supported under Grant Number [16/RC/3948] and co-funded under the European Regional Development Fund and by FutureNeuro industry partners, SFI Centre for Research Training in Genomics Data Science under Grant number [18/CRT/6214], and under the ‘Precision-ALS’ Spoke program of ‘FutureNeuro’ [20/SP/8953]. The authors also acknowledge Maria Bilen for sample preparation of Opa1 deficient animals.

## SUPPLEMENTARY FIGURE LEGENDS

**Supplementary figure 1. Diagram of how the sensitivity, specificity, and precision tests have been applied on simulated database.** Each run was replicated 10 times and TP: true positives, TN: true negatives, FP: false positivies, and FN: false negatives were scored.

**Supplementary figure 2. Low speed performance in previous sncRNA pipelines.** sncRNAP performs faster than tsRNAsearch and other tools such as smrnaseq.

**Supplementary figure 3. sncRNAP confirmed the identification of Val[C/A]AC and identified other sncRNAs candidates in hepatitis patients. A)** Read length distribution in bp of sncRNA reads. The majority of reads are between 20-25 and 30-35 bp. **B)** Percentages of RNA categories (miRNA, rRNA, and tRNA) for each sample. **C)** Counts of sncRNA classes for all samples. **D)** Pie chart showing the percentage for differentially and non-differentially expressed sncRNAs. E) Barplot showing the number of DE ((log2FoldChange > 1 or log2FoldChange < -1) & padj < 0.005) genes for each sncRNA class. F) Counts of the identified DE tRNA and miRNA families.

**Supplementary figure 4. Volcano plots showing the top DE sncRNAs in hepatitis patients.**

**Supplementary figure 5. sncRNAP confirmed the identification of miR135b and identified other sncRNAs candidates in PDAC. A)** Read length distribution in bp of sncRNA reads. The majority of reads are between 20-25 and 30-35 bp. **B)** Percentages of RNA categories (miRNA, rRNA, and tRNA) for each sample. **C)** Counts of sncRNA classes for all samples. **D)** Pie chart showing the percentage for differentially and non-differentially expressed sncRNAs. **E)** Barplot showing the number of DE ((log2FoldChange > 1 or log2FoldChange < -1) & padj < 0.005) genes for each sncRNA class.

**Supplementary figure 6. Volcano plots showing the top DE sncRNAs in PDAC.**

